# Novel nanopore sequencing method for determining Human Papillomavirus integrations in tumors without the need for whole genome sequencing

**DOI:** 10.1101/2024.10.17.618842

**Authors:** Preetiparna Parida, Nivedita Mukherjee, Agastya Singh, Shirley Lewis, Krishna Sharan, Sandeep Mallya, Ashima Singh, Mahadev Rao, Daniel S. Higginson, Radhakrishnan Sabarinathan, Rama Rao Damerla

## Abstract

Human papillomaviral (HPV) integrations into host human genome, a key event in cervical carcinogenesis, are currently mapped through laborious and expensive sequencing methodologies. We developed and validated a novel library preparation strategy for nanopore sequencing to generate long targeted reads with HPV and human chimeric sequences. Using this strategy, we validated known HPV integrations in HeLa (HPV18) and SiHa (HPV16) cell lines. We also mapped integration sites in five HPV+ cervical cancer patients, which were confirmed by whole genome and Sanger sequencing. Our nanopore-based method provides a precise and efficient strategy to capture HPV integrations crucial for understanding tumorigenesis.

## Introduction

Human papillomavirus (HPV) infections are associated with oropharyngeal, anogenital, genital, head and neck, and cervical cancers (1). Cervical cancer is the fourth most common cancer in women worldwide, which according to GLOBOCAN 2022, accounted for more than 662,301 new cases reported and 348,874 deaths (2,3). Persistent HPV infection is the primary driver of more than 99% of cervical cancers (4), 70% of which are caused by high-risk HPV types 16 and 18 (5). HPV is a non-enveloped double-stranded DNA virus belonging to the *Papillomaviridae* family. It infects the basal cells of the cervical epithelium and replicates as these cells divide and differentiate to form the upper epithelial layers, eventually being shed from the top layer as progeny virions that initiate reinfection (6). HPV induces carcinogenesis by inhibiting the host tumor suppressor proteins, p53 and retinoblastoma protein (Rb) using the viral proteins-E6 and E7 respectively (7). Integration of the HPV genome into a host chromosome is also a crucial step in HPV-induced carcinogenesis and accompanies the formation of invasive cervical cancer that breaks through the basement membrane. Integration sites are more likely to be enriched at common fragile sites and open chromatin regions in the human genome (8–10). Large-scale genomic analyses have also uncovered hotspots of recurrent HPV integration with significant enrichment for sequences with microhomology between the human and HPV genomes at the breakpoints (11). Tumors with HPV integration show upregulation of viral oncogenes E6 and E7, as well as host genes at or near the site of integration (12). Recent studies have observed that genomic regions around HPV integrations are enriched with Gene Ontology (GO) terms for DNA repair, fueling the hypothesis that integration events in the vicinity of DNA repair and tumor suppressor genes could lead to increased genomic instability (13). Moreover, HPV integration has been linked with changes in host chromatin structure and, consequently, in gene regulation, including long-range interactions (10).

Numerous such studies highlighting the cis and trans effects of HPV integration in inducing and promoting cervical carcinogenesis have underscored the importance of HPV integration detection in cancer research, leading to a surge in developing technologies to study HPV integrations (9). The Cancer Genome Atlas characterized 228 primary cervical cancers and reported HPV integrations in all HPV18-positive samples and 76% of HPV16-positive samples (10). Campitelli *et al*. used these cell-viral junctions as a biomarker in circulating tumor DNA for analyzing a series of serum samples obtained from cervical cancer patients and reported that HPV integration can be used as a biomarker for the detection of minimal residual disease and subclinical relapse in HPV-associated cancers (14). Therefore, detecting HPV integration status and identifying the integration locus are crucial steps not only for understanding the role of HPV in cervical carcinogenesis, but also for clinical diagnosis and treatment monitoring in cervical cancer patients (9).

Various techniques have been employed to determine HPV integration sites in the human genome, such as detection of integrated papillomavirus sequences by ligation-mediated PCR (DIPS-PCR) (15), Restriction Site PCR (RS-PCR) (16), amplification of papillomavirus oncogene transcripts (APOT) and next-generation sequencing (NGS). RNA-based assays for detection of HPV integration sites have substantial biological and technological constraints because of the lower stability of mRNA in biopsy samples (15). Though assays like DIPS-PCR are dependent on DNA, the potential for detecting new integration sites is limited (9). Recently, NGS has been widely used for the genome-wide characterization of HPV integrations in cervical cancer (17). However, the method requires whole genome sequencing at high coverage (>30X) (18), and involves multiple steps (19). Considering the complexity of the procedure and the resulting data (19), this method, though accurate, is not suitable for clinical use (9). Further, the cost of reagents used for these assays can be a challenge for labs with less resources, space, and medium to low sample throughput (20). With lower reagent costs and a shorter sequencing time, portable Nanopore sequencers may provide greater flexibility in this aspect (20).

Nanopore sequencing technology is the fourth generation of sequencing technology. It is portable, provides real-time data and enables rapid sequencing in clinical settings (21). In this study, as a proof of concept, we developed a novel enrichment strategy to capture HPV-human integration breakpoints using nanopore sequencing. We validated this technology by mapping known HPV integrations from cell lines and patient samples. Furthermore, by demonstrating expression changes of cancer-associated genes near the integration sites in the patient samples, we highlight the functional significance of mapping HPV integration events.

## METHODS

### Cell Culture

Cervical cancer cell lines SiHa (HPV-16 positive), and HeLa (HPV-18 positive), procured from the National Centre for Cell Sciences (NCCS), Pune, India, were cultured in Dulbecco’s modified Eagle’s medium (DMEM) (HiMedia Laboratories Pvt. Ltd., Mumbai, India) supplemented with 10% fetal bovine serum (HiMedia Laboratories Pvt. Ltd., Mumbai, India) and 1% antibiotic/antimycotic solution (HiMedia Laboratories Pvt. Ltd., Mumbai, India).

### Patient Recruitment And Sample Collection

Ethical approval for the study was obtained from the Institutional Ethics Committee, Kasturba Medical College Manipal, Manipal Academy of Higher Education, Manipal, India Fresh tumor biopsies from five patients with cervical cancer, admitted to Kasturba Medical College, Manipal, were collected after obtaining informed consent and prior initiating cancer therapy. Patient clinical parameters like age, histological grade, comorbidity status, etc., were obtained from the medical records and summarized in Table 1. Biopsy samples were snap-frozen in liquid nitrogen or dry ice ethanol bath and stored at −80 °C until further use.

**Table 1:**
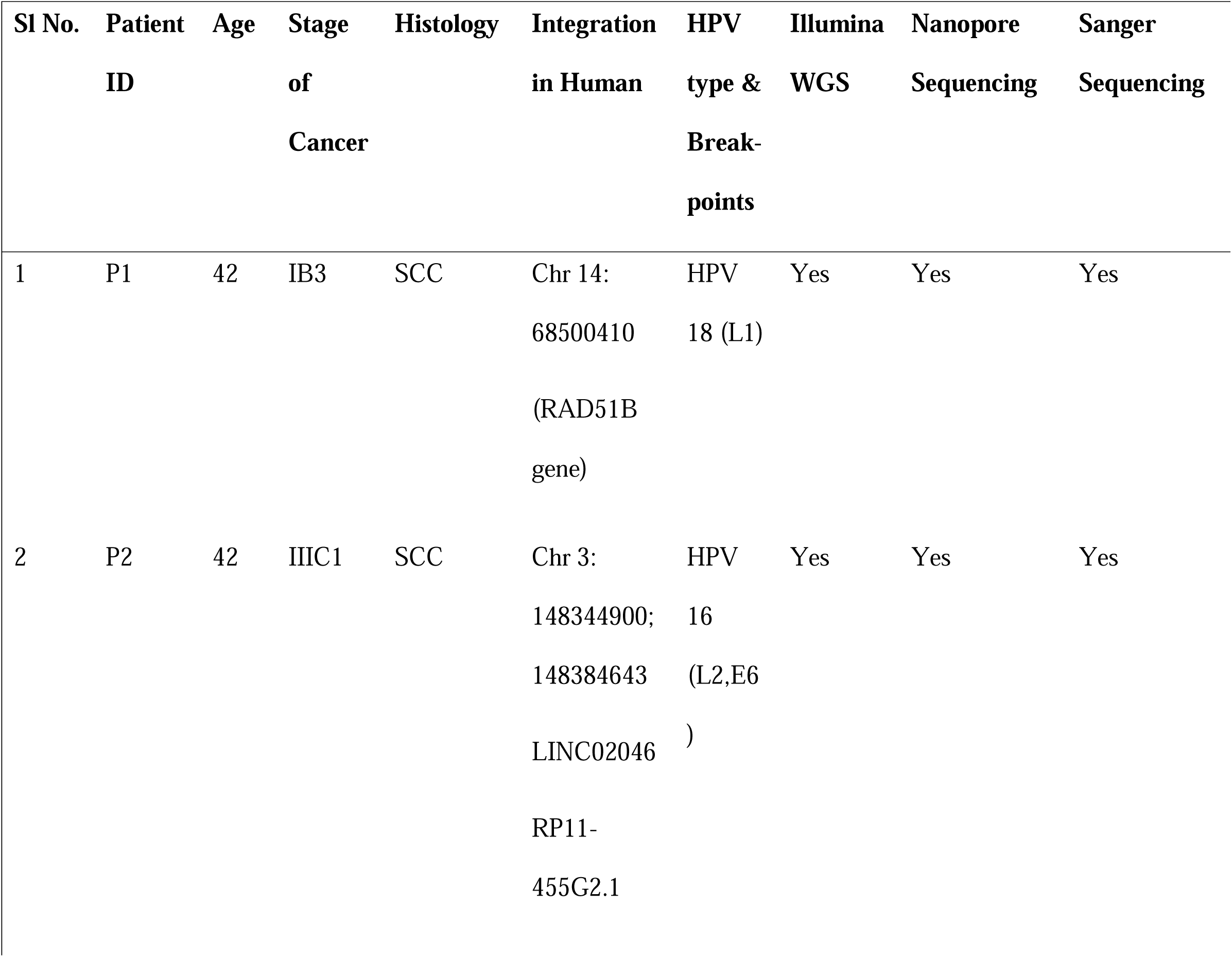
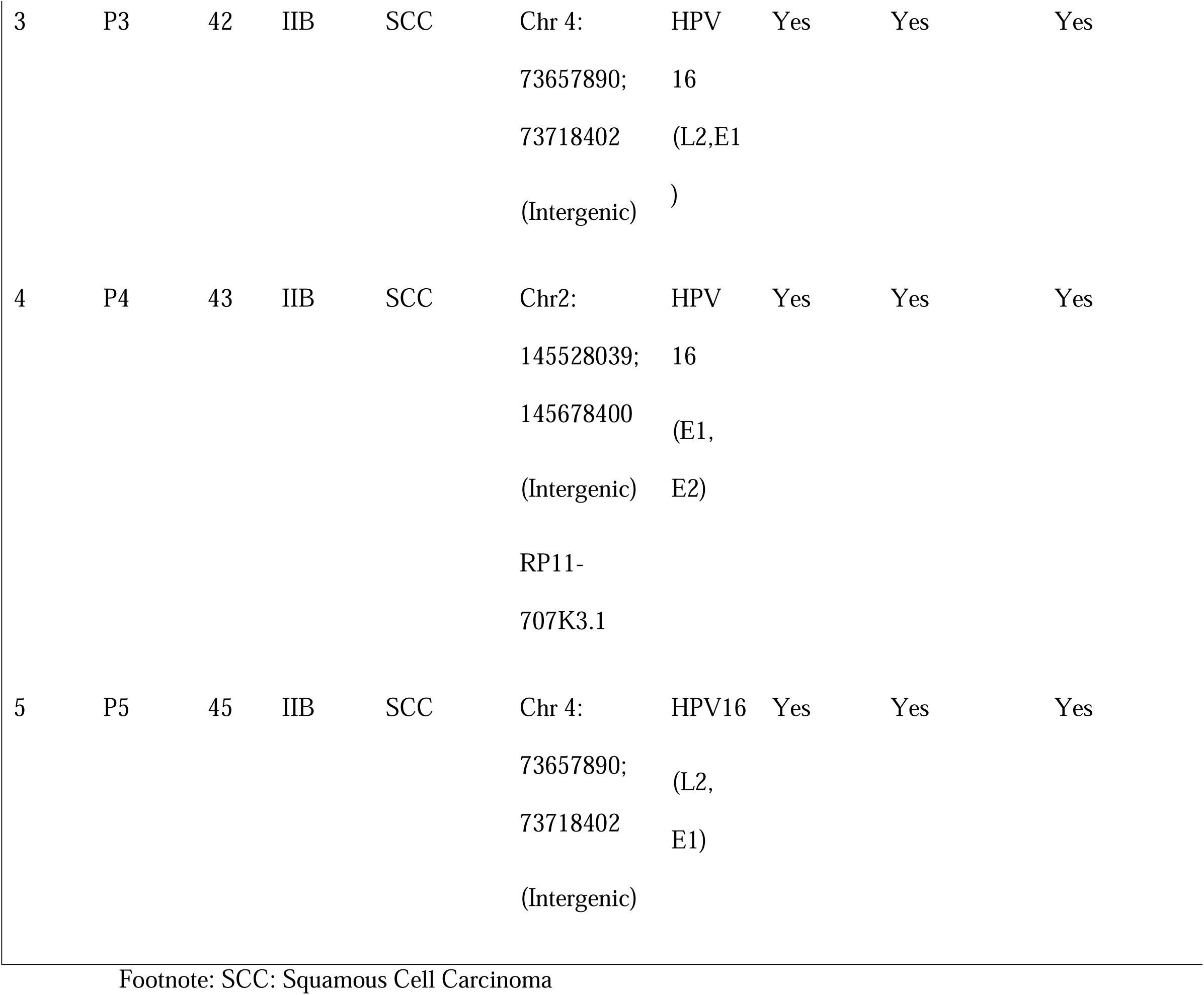
Demographic details of patients and comparison of breakpoint obtained by Illumina WGS and Nanopore sequencing.

### DNA Isolation, HPV Detection and Genotyping

DNA was extracted from tumor biopsy samples using a DNeasy Blood and Tissue kit (Qiagen, Hilden, Germany) following the manufacturer’s protocol. The DNA concentration of each sample was measured using Qubit® fluorometer 3.0 (Thermo Fisher Scientific Inc., USA) using the Qubit™ dsDNA BR Assay Kit. HPV detection by PCR was performed using HPV universal primers GP5+ and GP6+. PCR reaction comprised of 1x GoTaq® Green Master Mix (Promega Corp., WI, USA), 0.5 μM of GP5+ and GP6+ primers and 50 ng of genomic DNA. Thermal cycling was performed using SimpliAmp™ Thermal Cycler (Thermo Fisher Scientific Inc., USA) with the following conditions: Initial denaturation at 95°C for 2 min, 95°C for 30 s, 51°C for 30 s, 72°C for 30s, 35 cycles and final extension at 72°C for 5 min for 1 cycle.

### Whole Genome Sequencing (WGS) and analysis

Genomic DNA from tumor samples was sequenced using the Illumina Novaseq/ NextSeq sequencer as per the manufacturer’s instructions (Illumina, San Diego, California), with 2 × 150-bp paired-end reads and a minimum coverage of approximately 30X. WGS was performed at Medgenome laboratories based in Bangalore, India. FASTQ files from the paired-end WGS were passed through quality control and adapter trimming using the Trim Galore wrapper over Cutadapt and FastQC (22). To detect viral integration, the trimmed FASTQ files for each sample were processed through the ViFi algorithm using hg38 as the human reference genome, along with HPV genomes (23). ViFi employs phylogenetic methods in conjunction with reference-based read mapping to accurately identify integration events, even for novel viral strains. This analysis revealed samples with viral integration, providing details on the viral strain, the human chromosome and positional range of integration, as well as the number and identities of read pairs supporting the integration. Supporting reads include pairs in which one read maps to the human reference genome and the other to a viral reference genome, as well as split reads—a type of chimeric read that spans the integration site, mapping partially to the human and viral reference genomes. Split read sequences were fetched from the BAM file outputs from ViFi and verified by visualizing the alignments in the Integrative Genomics Viewer (IGV). Precise integration points were then identified from the split read sequences through BLAST analysis against the human reference genome and the detected viral strain.

### Single primer PCR extension

We developed a novel single primer extension strategy for specifically targeting HPV fragments. The schematic of the single primer extension based nanopore library preparation is depicted in Figure 1. This method for detecting HPV-human breakpoints involves a pool of primers flanking which cover the entire HPV genome. Two sets of primer pools were designed which amplify specific HPV genomic regions in forward and reverse directions respectively. The elongation of a single primer can capture HPV fragments and adjacent human genomic sequences during primer extension. The homopolymer tailing (C-Tailing) step followed by the single primer extension produces double-stranded DNA fragments and inhibits potential PCR-generated artefacts caused by errors in polymerase incorporation. This method can also detect novel integration breakpoints, unlike the traditional HPV detection PCRs which can only amplify a certain number of integration sites based on specific genes of HPV (E1, E2 OR L1) (15). Our method can also be instrumental in assessing HPV integration sites in highly fragmented or damaged DNA such as formalin fixed paraffin embedded (FFPE) tissue DNA, as it does not require a predefined amplicon length, unlike conventional PCR methods. The final round of nested amplification using nanopore adapter specific primers makes the assay more specific by enriching the amplicons obtained in the first round of primer extension. The use of nanopore specific adapter helps in barcoding, thereby multiplexing samples, which further reduces the cost of sequencing. The use of this library preparation technique for nanopore sequencing reduces the need for whole genome sequencing for detecting HPV-human integration events while increasing the efficacy of detecting HPV-human hybrid fragments even at very low copies.

**Figure 1:**
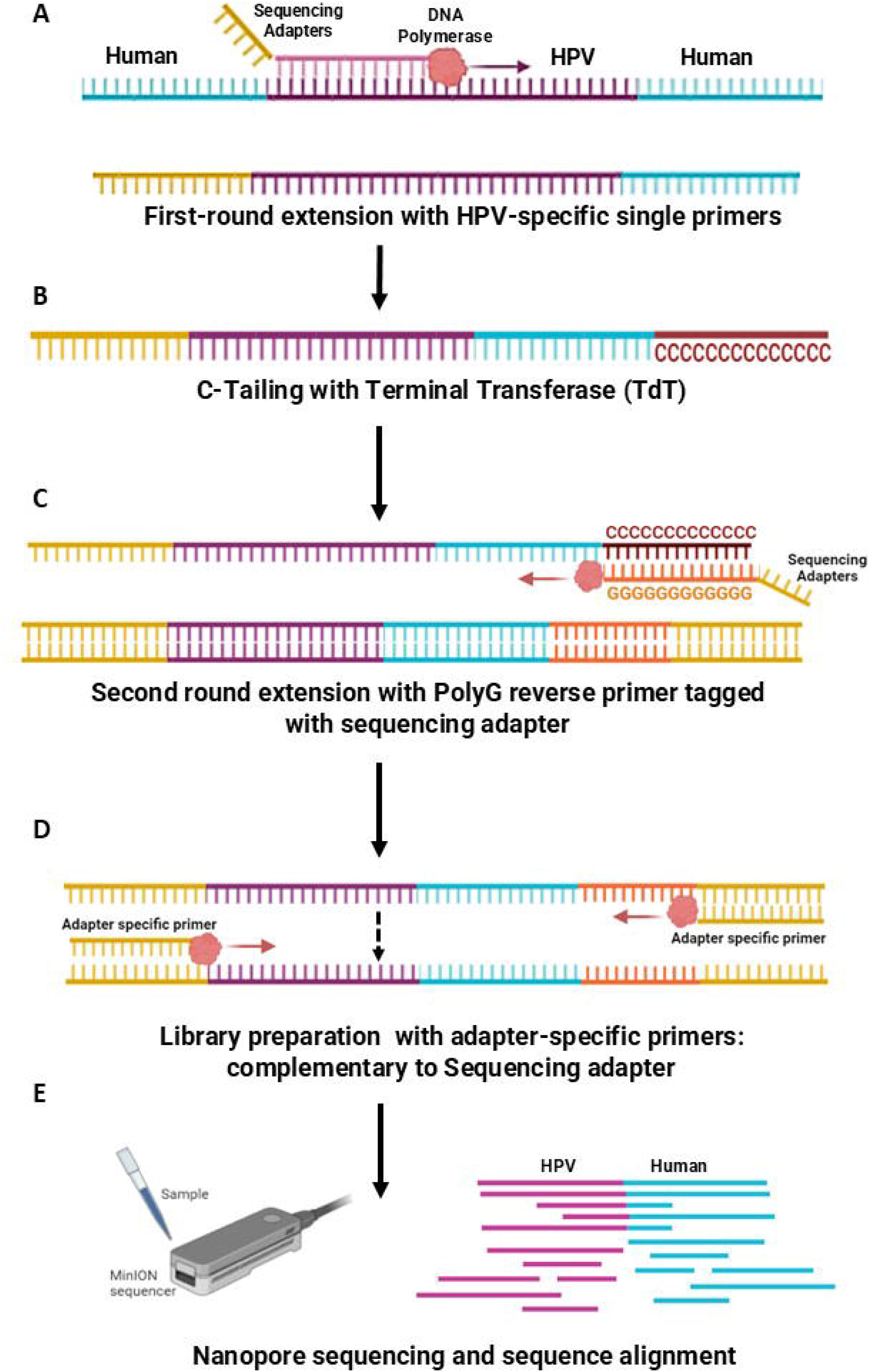
Schematic representation of Single primer extension method (A) Denaturation and hybridization of a single HPV-specific primer to the HPV sequence in the integrated region was followed by single Primer extension by a DNA polymerase; (B) HPV-specific single-stranded extended copies of the target DNA template molecules were then polyC-tailed using terminal transferase (TdT); (C) HPV-specific single primer-extended, polyC-tailed ssDNA molecules were then selectively amplified in second round extension with PolyG reverse primer tagged with sequencing adapter; (D) These products were used for library preparation for using adapter-specific primers: complementary to sequencing adapters. (E) Nanopore sequencing was performed and the sequences were analysed for identifying HPV-human integration sites. Created in BioRender. Genetics, M. (2024) BioRender.com/l69i229

### Primer Design

HPV16 and 18 whole genome references were obtained from the Papilloma Virus Episteme (PaVE) database (pave.niaid.nih.gov) (24). Primers were designed using PRIMER3 software such that they tile the whole HPV genome. These primers are listed in Supplementary Table 1. Tiled primer design was accomplished with 24 primers (10 in forward primer pool and 14 in reverse primer pool) each spanning an average of 500bp-1kb per genome of HPV16, and with 17 primers (9 in forward pool and 8 in reverse pool) each spanning an average of 1-1.5kb for HPV18.

### First round PCR Extension

First round of single primer extension was performed using 50 pmoles of HPV-specific primers with nanopore-specific adapters for barcoding, 500ng of genomic DNA extracted from the respective cell line or tumor samples, 10μl 5x PrimeSTAR GXL Buffer, 4μl dNTP mixture (2.5mM each), 1μl of PrimeSTAR GXL DNA Polymerase (1.25U/50μl) (Takara Bio, Cat# R050A). The PCR extension reaction was used to extend single-stranded long reads containing HPV sequences and the adjacent human genomic sequence. Thermal cycling was performed using SimpliAmp™ Thermal Cycler (Thermo Fisher Scientific Inc., USA) with the following conditions: 98°C for 10 s, annealing temperature (specific to the primer) for 30 s, and 68°C for 6 min for 50 cycles. Primer extensions were then size selected using 0.8x ratio of High Prep™ PCR Clean-up Beads (Magbio Genomics, USA) following the manufacturer’s instructions and eluted in 25 μL of nuclease and protease-free molecular biology grade water (HiMedia Ltd, Mumbai).

### C-Tailing

The purified amplification product (100 ng) was initially incubated for 5 min at 95°C in a water bath and chilled immediately on ice for 3 min. The nucleotide tailing reaction was then set up with 5 μl of 10X TdT buffer, 5 μl of 2.5 mM CoCl2, 1 μl of 100 pmol dCTP and 0.5 μl of terminal transferase (20 units/μl) (New England Biolabs), and the mixture was incubated for 1 hour at 37°C followed by heating for 10 min at 70°C. The reaction mixture was purified with paramagnetic beads and eluted in 10 μl of distilled water, and 5 μl was used for the second round PCR.

### Second round PCR extension

The second round PCR included a polyG reverse primer with nanopore-specific adapters for barcoding, 5μl of C-tailed products of first round of single primer extension, 10μl 5X PrimeSTAR GXL Buffer, 4μl dNTP mixture (2.5mM each), 1μl of PrimeSTAR GXL DNA Polymerase (1.25U/50μl) (Takara Bio, Cat# R050A). This PCR was performed under the following conditions: 98°C for 30 s, annealing temperature (specific to the primer) for 30 s and 68°C for 6-10 min for 40 cycles followed by a 10-min incubation at 68°C. These PCR products were purified using High Prep™ PCR Clean-up System (Magbio Genomics) and eluted in 25μl.

### Final PCR enrichment

The final round of PCR amplification included primers specific to adapter sequences, 10 μl purified product from the second round PCR, 10μl 5x PrimeSTAR GXL Buffer, 4μl dNTP mixture (2.5mM each), 1μl of PrimeSTAR GXL DNA Polymerase (1.25U/50μl) (Takara Bio, Cat# R050A). This PCR was performed with the following conditions: 98°C for 30 s, annealing temperature (specific to the primer) for 30 s, and 68°C for 10 min for 40 cycles followed by a 10 min incubation at 68°C. PCR products were purified using High Prep™ PCR Clean-up System (Magbio Genomics) and eluted in 25μl TE buffer. These purified products were processed for PCR barcoding for nanopore sequencing according to the manufacturers’ instructions. The schematic representation of the single primer PCR extension workflow is shown in Figure 1. The list of primers used in this assay is attached in Supplementary Table 1.

### PCR barcoding

PCR barcoding was performed using the barcodes provided in the PCR Barcoding Expansion 1– 12 kit (EXP-PBC001, Oxford Nanopore Technologies, Oxford, UK). One barcode was used per sample. The barcoding PCR reaction contained 2 μl PCR Barcode (one of BC01-BC12, at 10 μM), 500ng of purified PCR amplification product, 10μl of 5X PrimeSTAR GXL Buffer, 4μl of dNTP mixture (2.5mM each), 1μl of PrimeSTAR GXL DNA Polymerase (1.25U/50μl) (Takara Bio, Cat# R050A) and nuclease-free water up to 50 μL. The cycling conditions used for barcoding PCR consisted of 35 cycles with Initial denaturation 1 cycle of 95 °C for 3 mins, 35 cycles of denaturation at 95 °C for 30 seconds, annealing at 62 °C for 30 seconds, extension at 65 °C for 6-10 minutes, final extension 1 cycle at 65 °C for 10 minutes and hold at 4 °C. PCR barcoded products were purified using magnetic beads and the concentrations were measured using Qubit® fluorometer 3.0 (Thermo Fisher Scientific Inc.). These products were pooled in equimolar concentrations and 1 μg of pooled barcoded libraries was diluted in 47 μL of nuclease-free water for nanopore sequencing.

### Nanopore Sequencing

The barcoded library was then end-prepped, ligated with adaptors, and cleaned up for sequencing using the ONT sequencing ligation kit SQK-LSK109 kit (Oxford Nanopore Technologies). Qubit® fluorometer 3.0 (Thermo Fisher Scientific Inc.) was used to determine the concentration of the generated library. 50 fmol of the prepared library was loaded onto a R9.3 flow cell (ONT), after priming the flow cells with 800 µL of priming mix (30 µL Flush Tether to 1.17 mL of Flush Buffer). To prepare the library for loading, 11 µL (50 fmol) of the prepared library was mixed with 34 µL Sequencing Buffer, 25.5 µL pre-mixed loading beads, 4.5 µL nuclease-free water. 200 µL of priming mix was added to the priming port again avoiding the introduction of air bubbles. Finally, 75 µL of the sample mix was added to the flow cell SpotON sample port of the R9.3 flow cells (Oxford Nanopore Technologies) on the MinION in a dropwise manner. The samples were sequenced on the MinION Sequencer for 3 hours. Fastq files obtained from the nanopore sequencing run were analysed using the nfcore/nanoseq pipeline version 3.1.0 on Nextflow version 22.10.4. The raw fastq files were cleaned using NanoLyse and QC was done using FastQC. The processed reads were mapped to a custom genome consisting of human genome (hg38), HPV16, HPV18 and HPV31 sequences using minimap2 aligner. The chimeric soft clip reads from the output .bam files were extracted for further analysis. Circos plots were generated using the online tool shinyCircos (https://venyao.xyz/shinycircos/) (25).

### Viral Integration Detection Using Nanopore Reads

The reads obtained from Nanopore sequencing were aligned to a custom reference genome file using NGMLR-a long read aligner. NGMLR (https://github.com/philres/ngmlr) is designed to quickly and correctly align reads spanning complex structural variations and uses a convex gap cost model to compute precise alignments. It penalizes gap extensions for longer gaps less than for shorter ones thus accounting for large structural variants. The custom reference genome was created by concatenating hg38 and HPV reference genomes (HPV16 & HPV18). The aligned bam files (NGMLR output) were then used as input to Sniffles2 (https://github.com/fritzsedlazeck/Sniffles), a structural variation caller for long-reads. Sniffles2 outputs all types of detected structural variants (deletions, insertions, inversions and breakends). Only the breakends spanning human and viral contigs (with minimum five reads support) were considered as viral integration points.

### Sanger Validation

To validate the HPV integration sites in the human genome detected by nanopore sequencing, primers were designed, derived from the human genome at the potential site of integration, and the other against HPV sequences suspected of being near the site of integration within the HPV genome. Both primers were designed 500 bp away from the integration sites detected from the nanopore sequencing results. The PCR reaction mix was prepared in a total volume of 25 μL containing 12.5 μL 2X GoTaq Green Master Mix, 0.5 μL of each forward and reverse primer (10 mM), and 10.5 μL nuclease-free water. 1 μL (30 ng) genomic DNA solution was used as a template. The PCR conditions were as follows: 5 min at 95◦C; 35 cycles of 30 s at 94◦C; 60 s at 50–60◦C for annealing; and 60 s at 72◦C; followed by 72◦C for 1 min. The PCR products were run on 1.5 % TAE (Tris base, acetic acid and EDTA) agarose gel, the bands were excised and purified using FavorPrep Gel/PCR purification mini kit (Favorgen Biotech. Corp., Taiwan, Japan). Sequencing of the purified PCR products was performed using a BigDye® Terminator v3.1 Cycle Sequencing Kit (Applied Biosystems, USA). The list of primers used for validation is attached in Supplementary Table 2

### Functional validation of Impact of HPV integrations on the neighbouring genes

We wanted to explore the impact of the HPV integrations on the host gene expression in the context of topologically associating domains (TADs). For shortlisting target genes, cancer-related functions were determined from the IntOGen database (26) and existing literature. Promoter-enhancer interaction loop regions were obtained from the GeneHancer database (27). TAD boundaries of HeLa and NHEK cell lines were obtained from existing literature (28). All coordinates were obtained in hg38, or converted from hg19 to hg38 using UCSC LiftOver (https://pubmed.ncbi.nlm.nih.gov/16381938/). Total RNA was extracted from the five cervical cancer frozen tumor tissue and a control fibroblast cell line using TRIzol™ Reagent (Thermo Fisher, Carlsbad, CA, USA) according to the manufacturer’s protocol. 1 μg of total RNA was converted into cDNA using an iScript™ gDNA Clear cDNA Synthesis Kit (BioRad, Hercules, CA, USA) according to the manufacturer’s instructions. The reaction was incubated at 42°C for 30 minutes, followed by 85°C for 5 minutes to inactivate the reverse transcriptase. Primers were designed using primer3 software for qPCR. The list of primers used for qPCR is enlisted in Supplementary Table 3.

### Reverse Transcriptase-Quantitative Polymerase Chain Reaction Analysis of Selected Genes

We performed qPCR analysis for the panel of selected genes. To assess whether gene expression changes were specifically associated with HPV integration breakpoints and not due to general HPV infection, each patient was analyzed as a case, with the other four patients serving as controls. This intra-cohort comparative design ensured that differences in gene expression were linked to the integration event and not HPV infection. The 2^−ΔΔCt^ method was used for quantification and fold change for the target gene. RT-qPCR analysis was performed using TB Green Premix Ex Taq II (Tli RNase H Plus) (Takara Bio Inc., Tokyo, Japan; RR820A) on QuantStudio^TM^ 5 (Applied Biosystems, Waltham, MA, USA). The PCR cycling conditions were as follows: 95 °C for 1 min followed by 40 cycles of 95 °C for 5 s, annealing temperature (specific to the different genes) for 30s, and 72 °C for 30s. The values were first normalized to the internal reference gene GAPDH, followed by calculating relative expression to healthy control. Statistical significance was inferred using *t*-test, with a *p*-value < 0.05 considered statistically significant.

### Sequencing Cost Analysis

The cost per sample was calculated by considering flow cell costs, sequencing kits, wash kits, PCR reagents, and terminal transferase. It also includes all other sample preparation and purification reagents before sequencing. WGS was performed at Medgenome laboratories, India; hence, the price charged per sample for sequencing was considered.

## RESULTS

### Single primer extensions for detection of HPV integrations in HPV Positive Cell Lines

We developed a single primer extension method to capture HPV-human breakpoints by designing primers tiling the genomes of the most commonly occurring high-risk HPV subtypes, HPV16 and HPV18. The single primers allowed us to randomly extend the HPV genome allowing us to capture integrations provided the DNA sequence adjacent to the primer binding site is in the vicinity of a HPV-human integration junction. The addition of poly-C tail by the terminal transferase enzyme enables the binding of a reverse primer with poly-G resulting in a double-stranded molecule with nanopore adapters at either end. The reads obtained from the nanopore sequencing were in size ranging from 96 to 4656 base pair (with a median size of 228). On an average, the polyC and polyG tracts observed were 7 to 8 base pairs long. To establish and validate this assay, we used genomic DNA from HPV-positive cervical cancer cell lines, HeLa and SiHa, to map chimeric viral-human breakpoints. Our analysis detected all the previously reported viral-human breakpoints in these cell lines (with >100 reads supporting the integration).

We could accurately identify integration sites on chromosome 8 of the HPV18-positive HeLa cell line as reported previously using the DIPS PCR technique Luft *et al.* (15,29) (Figure 2A, Supplementary Figure 1A). Similarly, using this method, we effectively detected the integration sites on chromosome 13 in the HPV16-positive SiHa cell line as reported by Meissner (30) and Yu *et al*. (29). To validate our findings, we used the Sanger sequencing approach to ensure the accuracy of detection of viral breakpoints in these cell lines (Figure 2B, Supplementary Figure 1B)

**Figure 2:**
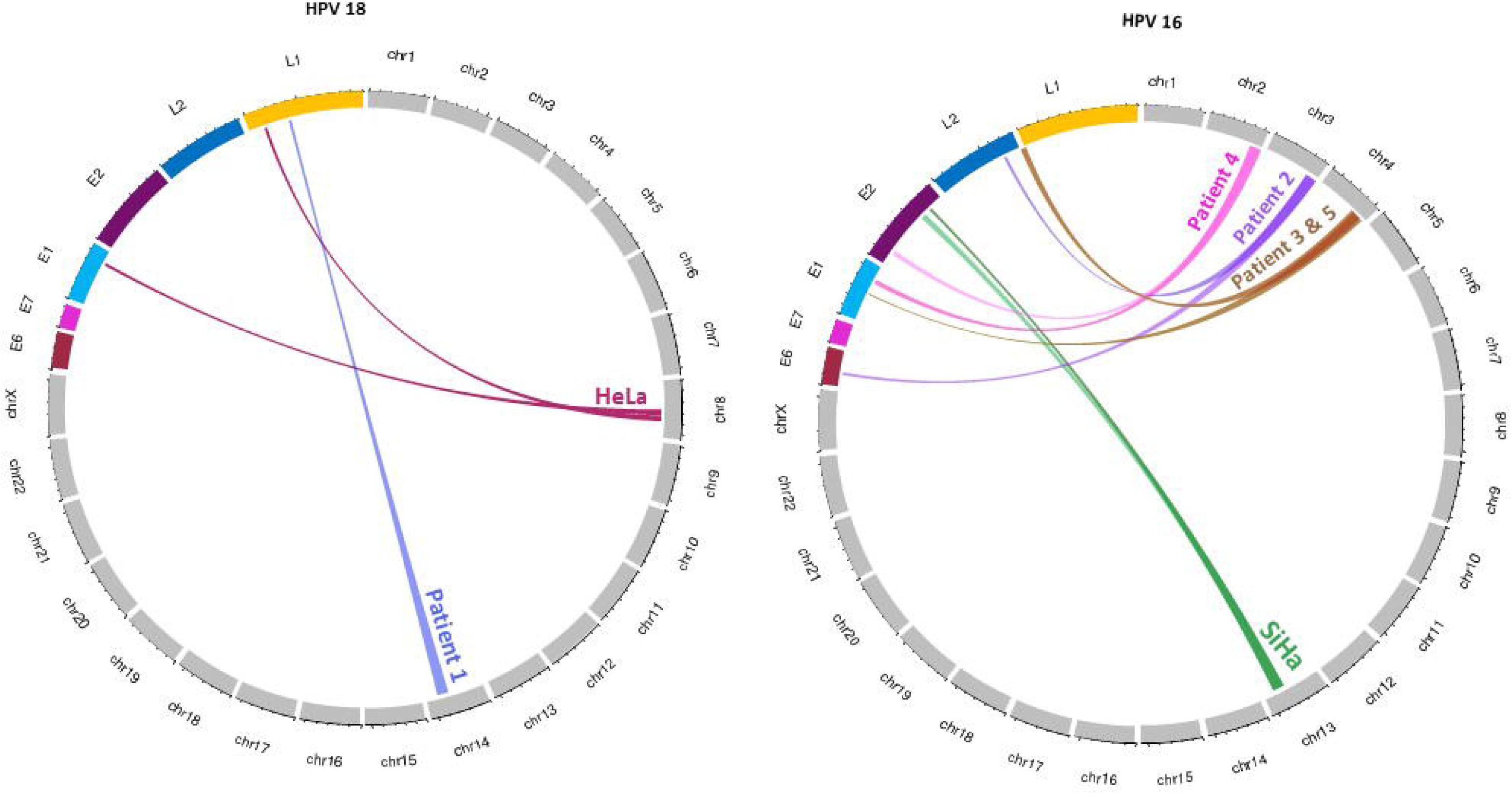
Circos plot illustrating the chimeras between the HPV genome (specific gene) with specific human chromosomes A. Illustrating chimera of HPV 18 with HeLa cell line (in Chromosome 8) and Patient P1 (in Chromosome 14); B. Illustrating chimera of HPV 16 with SiHa cell line (in Chromosome 13), Patient P2 (in Chromosome 3), Patient P3 (in Chromosome 4), Patient P4 (in Chromosome 2) and Patient P5 (in Chromosome 4).

**Figure 3:**
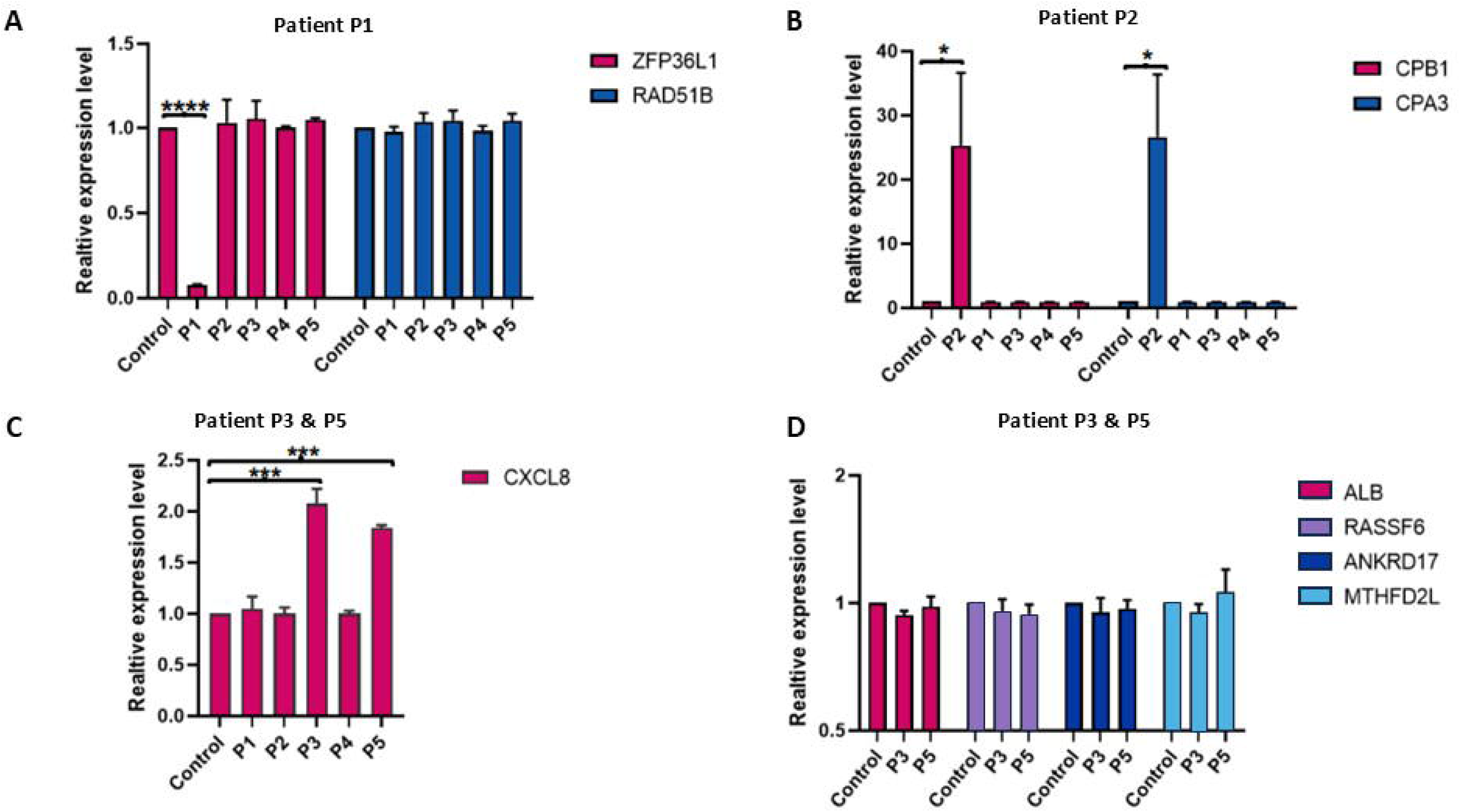
Relative expression of genes associated with HPV integration with respect to their TAD domains. A) Relative gene expression of *ZFP36L1* and *RAD51B*. B) Relative gene expression of *CPB1* and *CPA3*. C) Relative gene expression of *CXCL8*, D) Relative gene expression of *ALB, RASSF6, ANKRD17* and *MTHFD2L*.

### Characterization of HPV Integrations in tumor samples of cervical cancer patients

We used the single primer extension method for analyzing HPV integration sites in five HPV-positive cervical cancer patients (details summarized in Table 1). Patient P1 had an integration of the viral genome into intron 4 of the RAD51 paralog B (*RAD51B*) gene on chromosome 14. We could find only one viral breakpoint within this gene, and the integration resulted from the disruption of L1 gene of the virus. Patient P2, had an integration of HPV genome in a lncRNA gene LINC02046 on chromosome 3. Three patients P3, P5 and P4 had integration sites in intergenic regions of Chromosome 4, and Chromosome 2 respectively. Patients P3 and P5, had the exact same integration sites on chromosome 4. WGS was performed for the five patients using Illumina to validate the results of our analysis. Sequencing analysis from WGS demonstrated concordance with nanopore sequencing data (Table 1). We further validated these integration sites by Sanger sequencing (Supplementary Figure 2). The breakpoints obtained for the five patients through Illumina WGS and Nanopore sequencing are mentioned in Table 1 and Supplementary Figure 2.

HPV integration has been shown to disrupt host transcriptional activity by upregulating the expression of nearby genes through chromatin remodeling (31). This transcriptional disruption is mostly confined to genes present within the same TAD (Topologically Associated Domain)– functional units of 3D genome organization that harbor genes and regulatory regions and spatially confine gene interactions (32). To evaluate the influence of HPV integration on transcriptional activity in the five cervical cancer patients, we shortlisted target genes located within the same TAD as the integration sites (Supplementary Figure 3-6).

For patient P1, cancer-associated genes located within the same TAD were selected, including RAD51B, which overlaps with the integration site, and ZFP36L1. In patient P2, the HPV integration occurred near the 5’ TAD boundary. Hence, genes near the 3’ TAD boundary, potentially interacting with the integration site through TAD boundary interactions, were selected. This included CPB1 (within the TAD near its 3’ boundary) and CPA3 (outside the TAD but near its 3’ boundary). For patients P3 and P5, which had the same integration site, genes CXCL8, RASSF6, ANKRD17-DT and MTHFD2L were selected since the integration site fell within their promoter-enhancer integration loops. Additionally, the cancer-associated gene ALB, located within the same TAD, was also selected. No target genes were selected for P4, as the integration site was located outside any TAD, thus devoid of genes or regulatory regions.

In patient P1, *ZFP36L1* was downregulated compared to the control sample and the other four patients, while the expression of *RAD51B* did not exhibit any significant difference. Genes *CPA3* and *CPB1* were found to be significantly upregulated in patient P2 as compared to the control as well as the other four cervical cancer samples. Patients P3 and P5 shared the same integration breakpoint. Among the cluster of genes selected for P3 and P5, only the relative gene expression CXCL8 was upregulated in both patient P3 and P5 but was normal for the other 3 samples and control.

### Sequencing Cost

The cost per sample for WGS was $598.74 while the final cost of sequencing per sample on MinION using our novel library preparation technique was $27. The final cost was calculated assuming 10 samples per run and did not include labor charges. For calculating the per-run cost for purification, we assumed each sample volume to be 50µL, and in each run, the magnetic bead purification was performed five times. Each flow cell was washed and reused (4 runs per flow cell). When we included the reagent costs for single primer assay along with nanopore sequencing, our cost per-sample was $54.6. Detailed price analysis is included in Table 2.

**Table 2:** Cost analysis for MinION Nanopore sequencing using PCR enrichment method.

| Nanopore Sequencing | Cost (in US\$) | Total Units | Cost per run (in US \$) |
| --- | --- | --- | --- |
| PCR | 148.9 | 250 | 0.6 |
| C-tailing (Terminal<br>Transferase NEB) | 213.65 | 500 | 0.4 |
| Magnetic beads for<br>purification HighPrep PCR | 123.77 | 125 (40ul per<br>reaction) | 5.0 (5 times per run) |
| Minion Flowcell R9.4.1 FLO-MIN106D | 1,081.16 | 4 runs per flow<br>cell | 270.3 |
| Ligation Sequencing and<br>adapter ligation (SQK-LSK109 kit) | 907.46 | 6 | 151.2 |
| Barcoding (PCR Barcoding<br>Expansion 1-12) | 454.74 | 6 | 75.8 |
| Flowcell Priming (EXP-FLP002) | 155.73 | 6 | 26.0 |
| Flowcell Wash (EXP-WSH004) | 99 | 6 | 16.5 |
| Total Cost |  |  | 545.8 |
| Cost per sample |  |  | **54.6** |

## DISCUSSION

Nanopore has emerged as an important next generation sequencing tool since its release in 2014 (18). In this study, we developed a novel targeted sequencing approach which combines single primer extension enrichment followed by nanopore sequencing to precisely identify HPV integration sites in HPV-positive cell lines and tumor genomic DNA from cervical cancer patient samples. Whole genome sequencing on the Nanopore sequencing platform has been previously used for studying viral integration due to its ability to generate long reads, enabling comprehensive genome-wide analyses (9). Increasing the depth of sequencing by enriching the viral genome using a PCR can help in attaining improved accuracy (21). The findings of our study highlight the effectiveness of our unique targeted nanopore based amplicon sequencing in accurately identifying integration sites within known chromosomal regions of HPV-positive cell lines (HeLa in chr8 and SiHa in chr13). Our results conformed previously identified integration breakpoints in these cell lines (15,16).

We further evaluated the utility of our single primer based targeted nanopore sequencing by comparing our results with Illumina Whole Genome Sequencing (WGS). Our analysis of 5 cervical cancer patient samples reported concordance with Illumina WGS. In our study, one patient (Patient P1) had an integration within intron 4 of the RAD51B gene located on chromosome 14. RAD51B is an important DNA double-strand break repair protein, and its loss of function is reported in uterine leiomyoma and breast cancer (33). It has also been previously reported to be one of the hotspots for HPV integration which may lead to dysregulation of Homologous Recombination Repair (HRR), causing genomic instability—a hallmark of cancer (34). We also observed that this integration led to the downregulation of a neighboring gene within the same TAD, Zinc finger protein 36 (*ZFP36L1*), which acts as a tumor suppressor. *ZFP36L1* is often downregulated in several patient cohorts of bladder and breast cancers and its reduced expression is associated with worse survival in patients with breast cancer (35). Loss of *ZFP36L1* has also been reported to promote epithelial-mesenchymal transition in hepatocellular carcinoma (36).

We also found an HPV integration site in a lncRNA gene LINC02046 in Patient P2. The sequencing data for the other three patients revealed integrations predominantly within intergenic regions, which is consistent with patterns reported in previous literature (11,29,37). This integration shared the TAD boundaries with two genes, *CPB1* and *CPA3*, which were found to be upregulated in the patient P2. It has been reported that the overexpression of *CPB1* in ductal carcinoma in situ (DCIS), which is an early-stage breast cancer, can lead to the progression into invasive breast cancer in patients (38). Also, gene *CPA3*, currently known as *CPA4* (39) is associated with pancreatic cancer progression, and was observed to be overexpressed in pancreatic cancer patients when compared to healthy controls (40).

Integration sites in the HPV-positive cell lines HeLa and SiHa were also observed in the intergenic regions of Chr 8 (Chr8q24.21) (11,29) and Chr13 (between the KLF5 and LINC00392 genes) (37). As hypothesized by Yang *et al.,* intergenic HPV integrations might serve as a defense mechanism of the virus which keeps the host cell viable as integration into human non-exonic regions would probably not cause a significant loss of function of any important gene (9). Interestingly, two patients in our cohort (P3 and P5) had the same intergenic integration site and breakpoint patterns. We observed that this integration break point fell within the promotor enhancer integration loops of genes in that TAD including *CXCL8, RASSF6, ANKRD17* and *MTHFD2L*. *CXCL8* was overexpressed in these two patients P3 and P5 as compared to the other three patients and control. None of the other genes, nor ALB, a cancer-associated gene present in the same TAD, showed significant changes in the expression. *CXCL8* is a known chemokine which plays major role in the proliferation, invasion, and migration of cancer (41).This gene is also reported to promote angiogenesis in breast cancer (42) and the overexpression of *CXCL8* is associated with increased risk of cancer and poorer prognosis in patients with colorectal cancer (43). This integration breakpoint can be further analyzed in a larger cohort of patients for its potential as a biomarker or to be considered as a hotspot.

We believe our method has many advantages over both Ilumina and Nanopore WGS. We developed a targeted method for the detection of HPV integration sites which is over 20 times more cost-effective than WGS and minimizing sequencing data volume. With our novel enrichment method, we could also enable deeper analysis at the specific region of interest (HPV-human Integration breakpoint) which helps to avoid false positive interpretation of sequencing data. Also, this strategic combination not only increased our sensitivity in detecting integration breakpoints but also substantially reduced background noise, enabling us to pinpoint specific integration loci within the genome.

The accurate identification and characterization of HPV integration breakpoints holds immense clinical implications, especially in understanding the molecular mechanisms underlying HPV-associated carcinogenesis (9). Our study highlights the potential of nanopore sequencing, complemented by the single primer extension enrichment method, as a useful tool for this purpose. Besides the reduced cost and amount of sequencing data, a simpler analysis pipeline makes HPV integration detection faster, more efficient, and easily adoptable, which is crucial for clinical applications. Larger cohorts may be explored in future studies to identify new integration sites, potentially paving the way for the development of prognostic biomarkers or targeted therapeutic approaches for HPV-associated cancers. Our method demonstrates the superiority of targeted nanopore sequencing in identifying HPV integrations compared to Illumina WGS, emphasizing its precision, comprehensive coverage, and compatibility with a novel enrichment method.

## CONCLUSION

Our study demonstrates the efficacy of a targeted nanopore sequencing approach combined with a novel enrichment method for precise and efficient detection of HPV integrations in genomes of cervical cancer patients. This method has an advantage over the traditional Illumina WGS in terms of cost-effectiveness, and data analysis efficiency. This method will contribute significantly to the understanding of the role of HPV integration in cervical carcinogenesis by accurately identifying integration breakpoints. Our novel technique could be easily applied to studying other viral integrations like retroviral, Adenovirus associated and Human Herpes-viral integration events. In addition to mapping viral integrations, this method could also be a crucial and a cost effective solution for detecting gene fusions in several cancers. Larger cohort studies are further required to explore the clinical significance of this method and translate it to diagnostics or companion diagnostic solutions.

## AUTHORS’ CONTRIBUTIONS

PP and RD conceived the project. PP, NM, RS and RD designed the experiments and wrote the manuscript with inputs from all authors. PP and AS2 performed the experiments along with nanopore sequencing. NM, AS1, SM and RS performed nanopore and WGS data analysis. SL and KS assisted in sample characterization and clinical data compilation. DH and MR helped with important inputs to the study design and critical review of the manuscript. RD and RS supervised the study. All authors reviewed and approved the submitted version of the manuscript.

## ACKNOWLEDGMENTS

PP would like to acknowledge the support from Senior Research Fellowship (SRF-Direct) (CSIRAWARD/SRF-DIRECT2024/15079) awarded by Council for Scientific and Industrial Research (CSIR), Ministry of Science & Technology, Government of India. NM would like to acknowledge support from NCBS-TIFR and the Shyama Prasad Mukherjee Fellowship (SPMF) awarded by Council for Scientific and Industrial Research (CSIR), Ministry of Science & Technology, Government of India. RD would like to acknowledge faculty seed money support from Manipal Academy of Higher Education, Manipal, Karnataka, India. RD and MR would like to acknowledge the support and fruitful discussions through the Global Cancer Consortium (https://glocacon.org/).

## DATA AVAILABILITY STATEMENT

The data that support the findings of this study are available from the corresponding author upon reasonable request.

## ETHICS STATEMENT

Ethical approval for the study was obtained from the Institutional Ethical Committee, Manipal Academy of Higher Education (IEC No: 774/2019). This study was also registered as an observational study with Clinical Trials Registry – India with the number CTRI/2020/01/022862.

---

**Supplementary Table 1:**
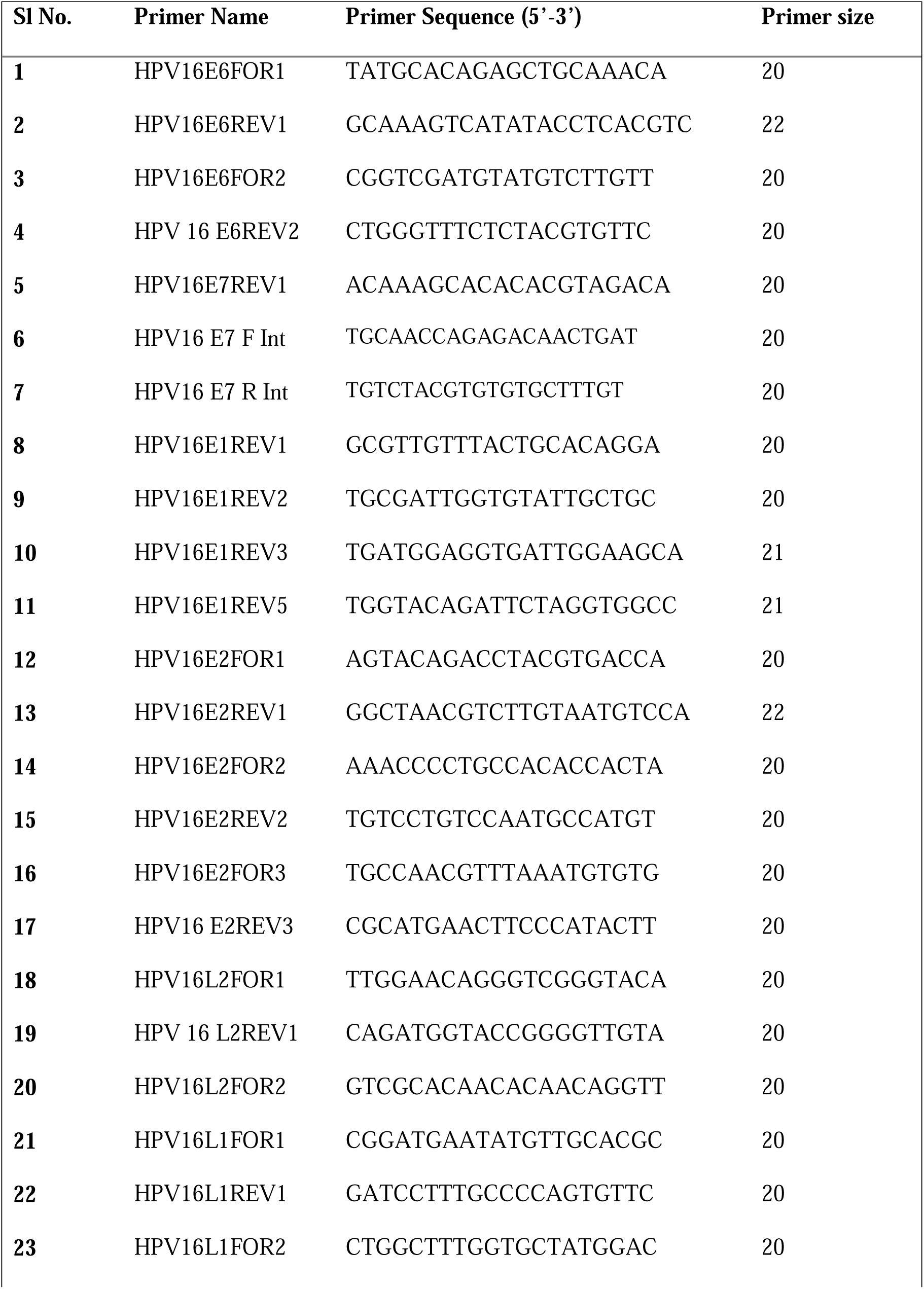
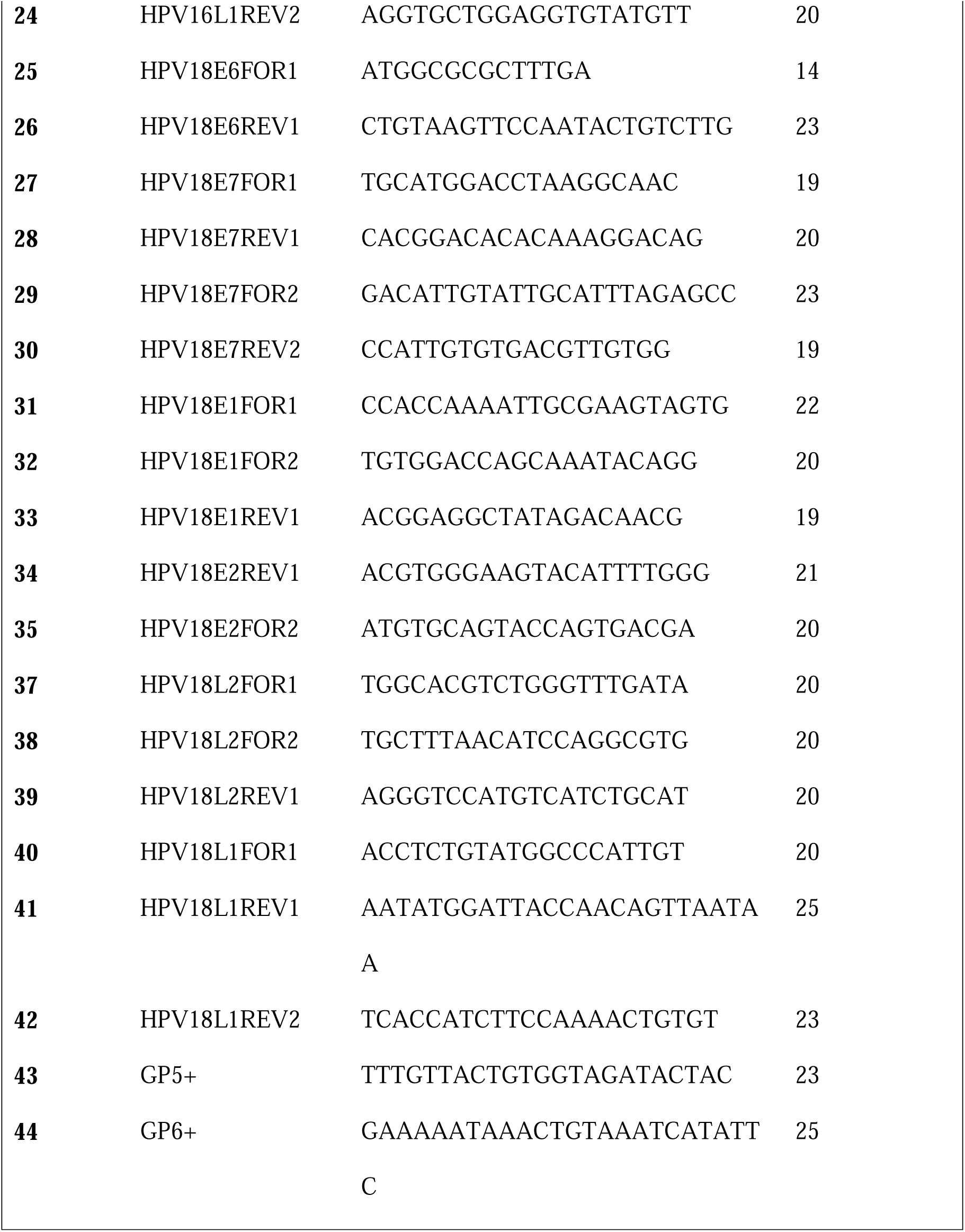
Primers used for Single Primer Enrichment.

**Supplementary Table 2:**
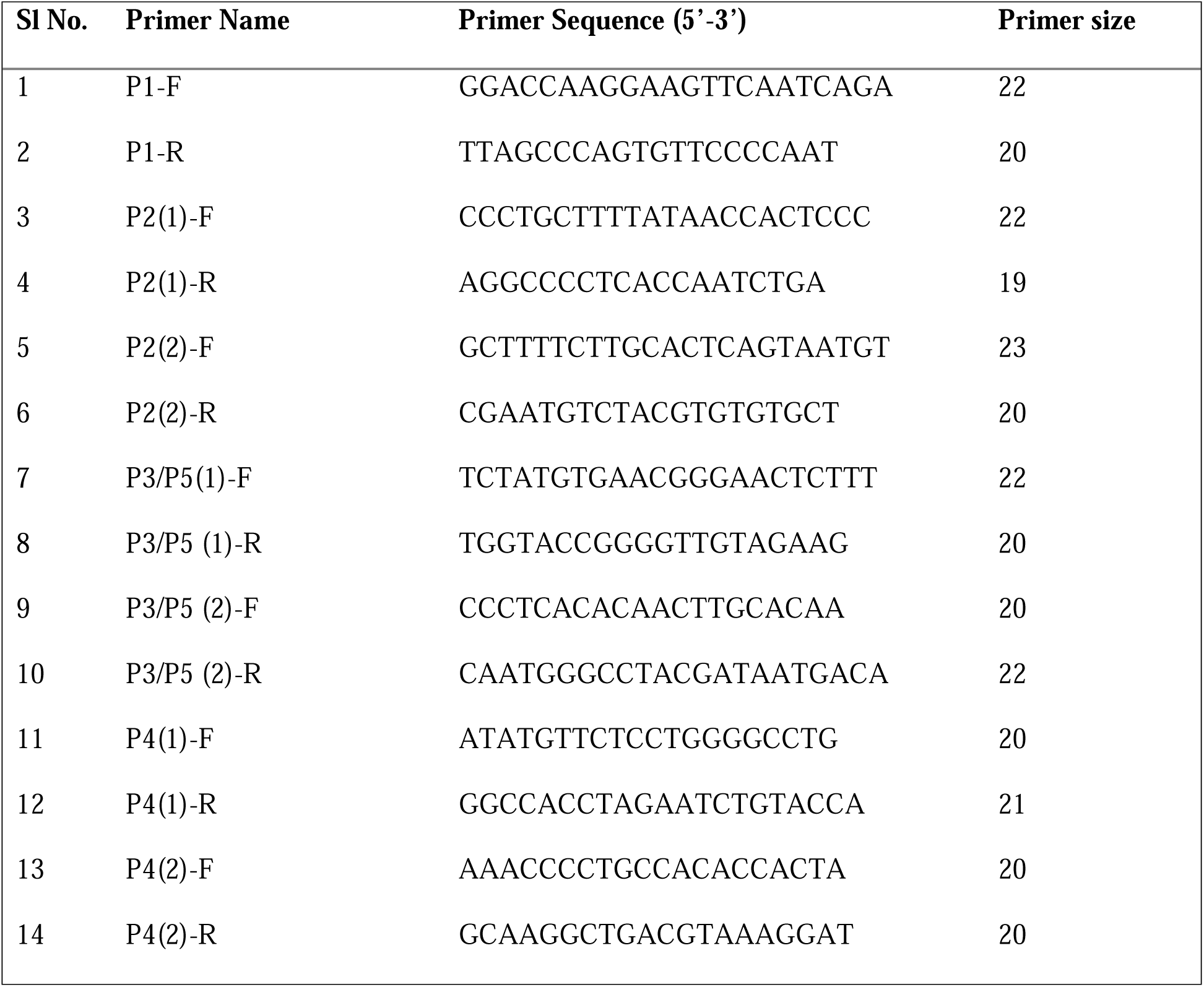
List of primers used for validation of breakpoints in cervical cancer samples.

**Supplementary Table 3:**
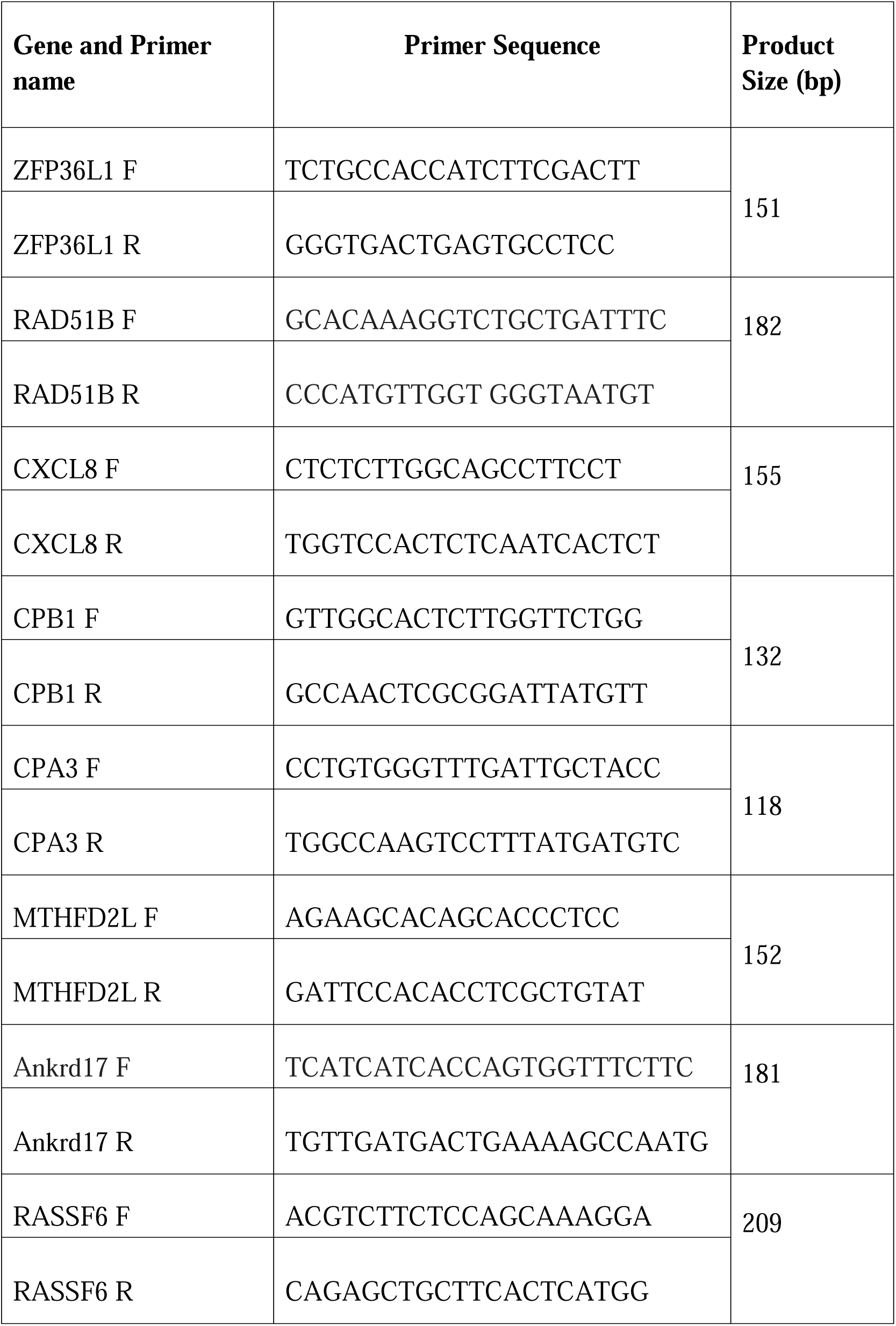
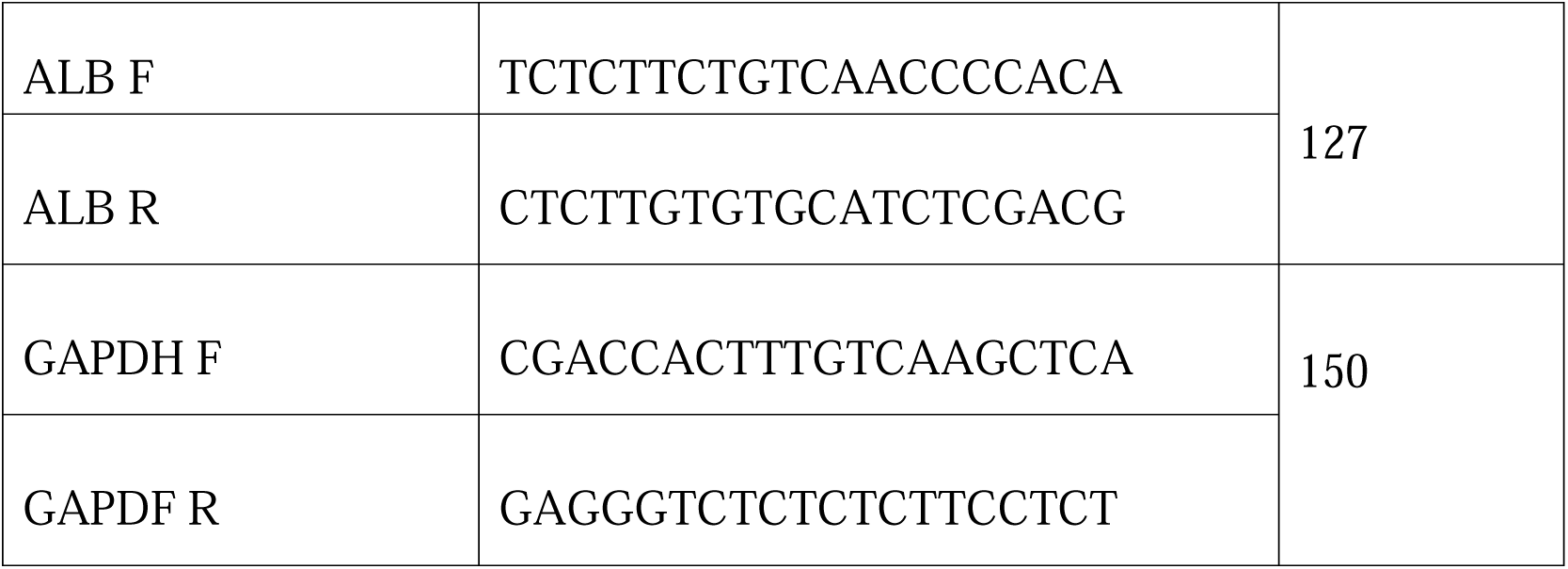
List of primers used for qRTPCR.

