## Supplementary figures and images for "Novel nanopore sequencing method for determining Human Papillomavirus integrations in tumors without the need for whole genome sequencing"

### Supplementary Figure 1

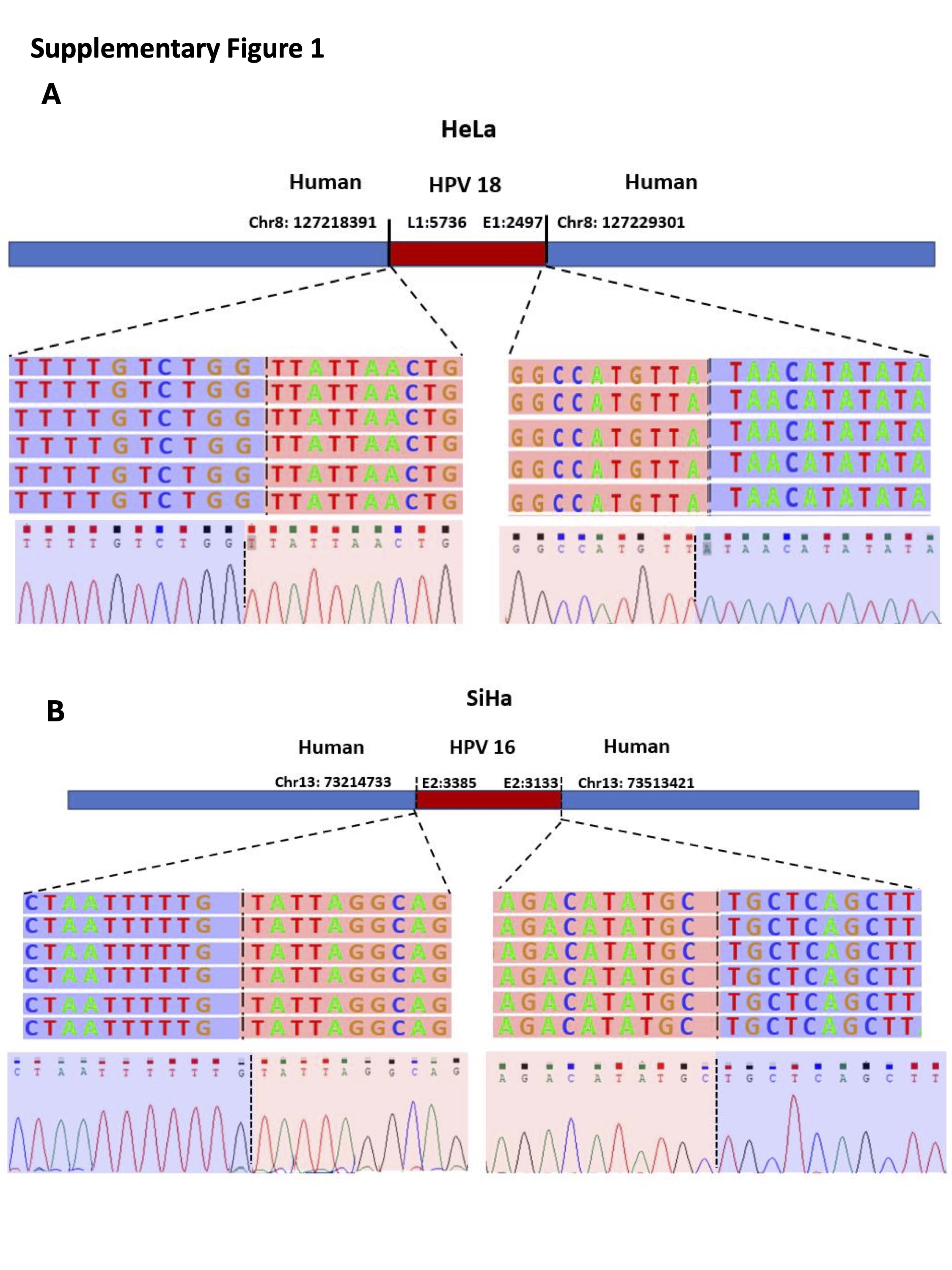

### Supplementary Figure 2

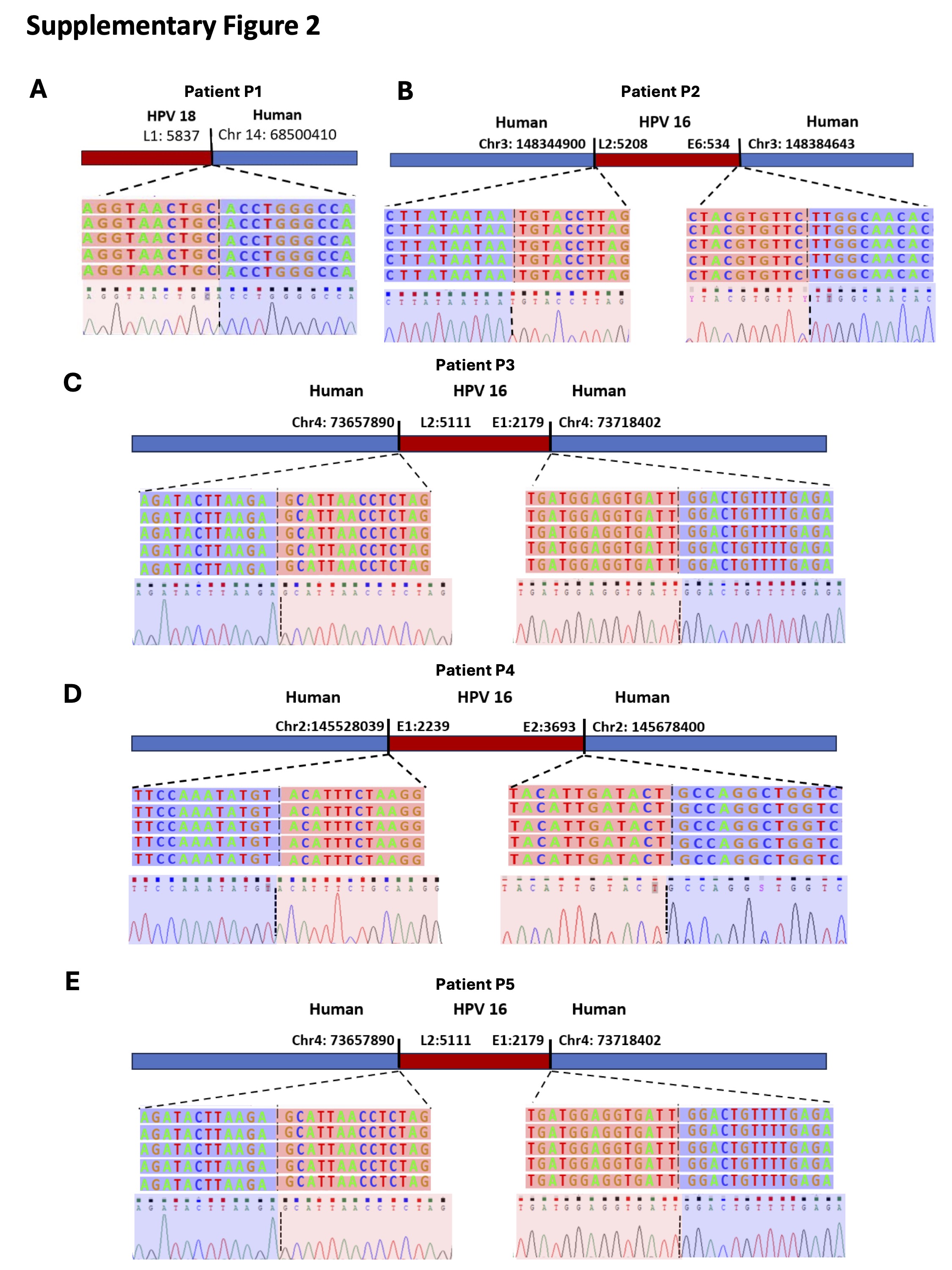

### Supplementary Figure 3

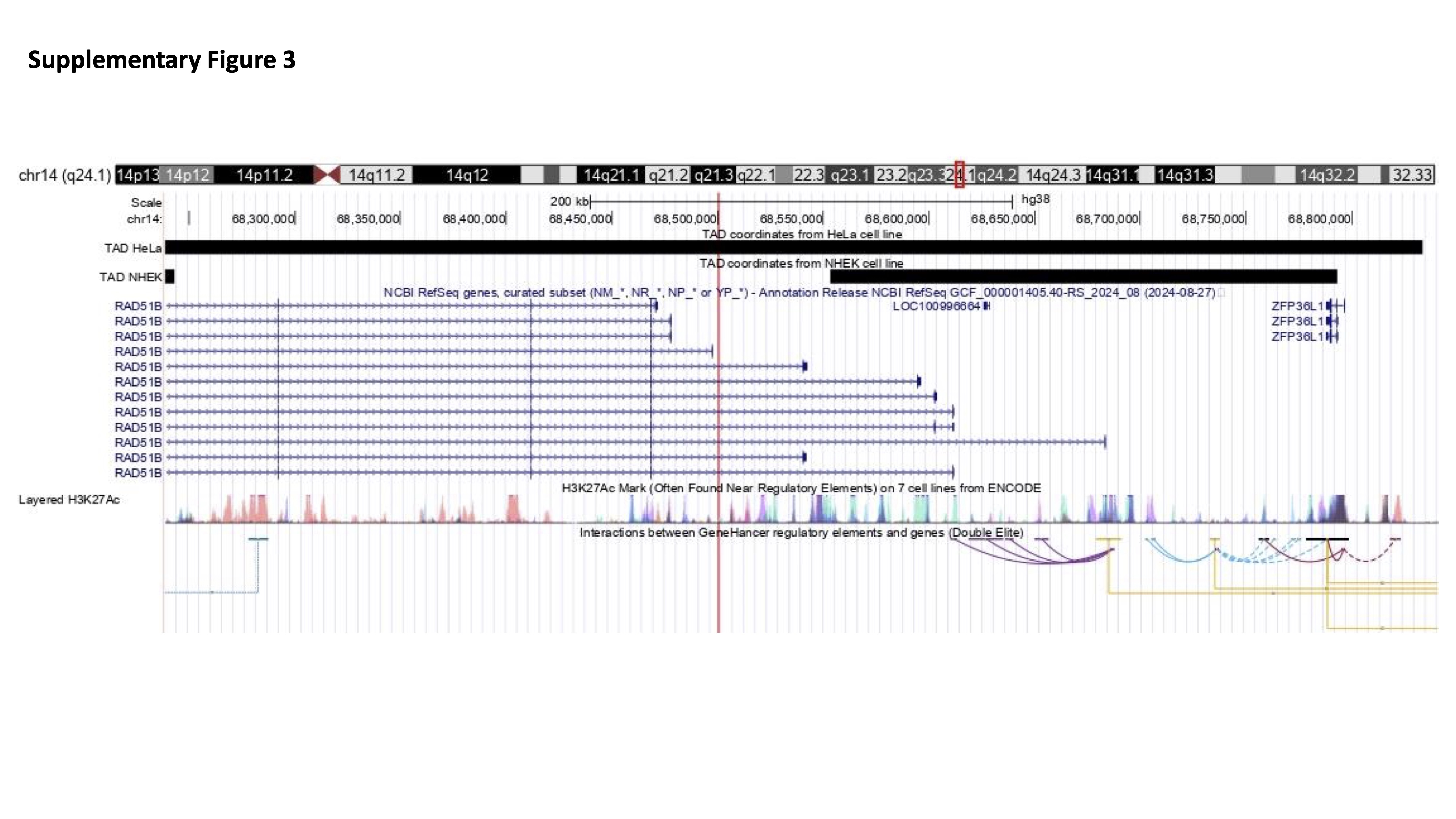

### Supplementary Figure 4

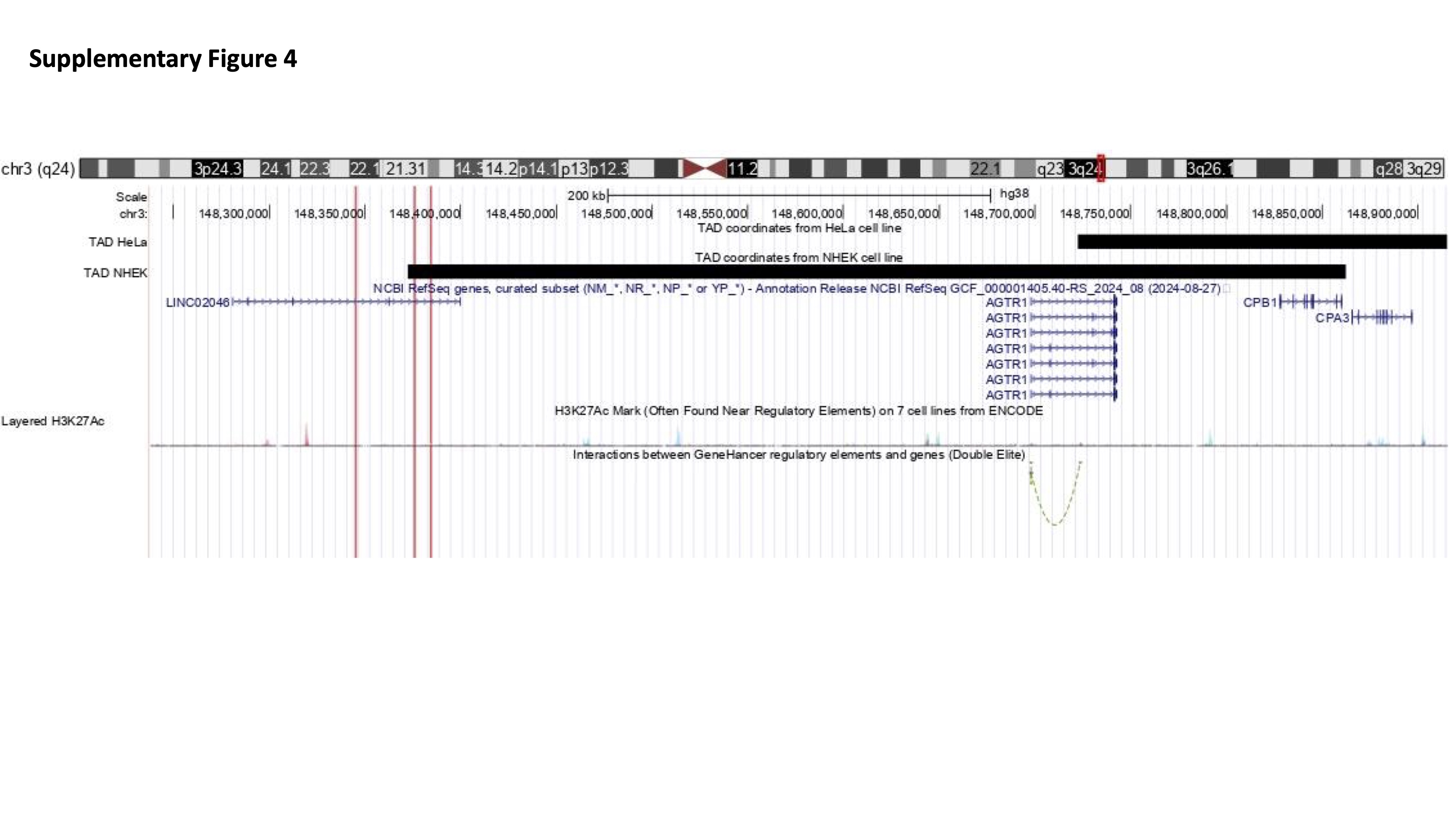

### Supplementary Figure 5

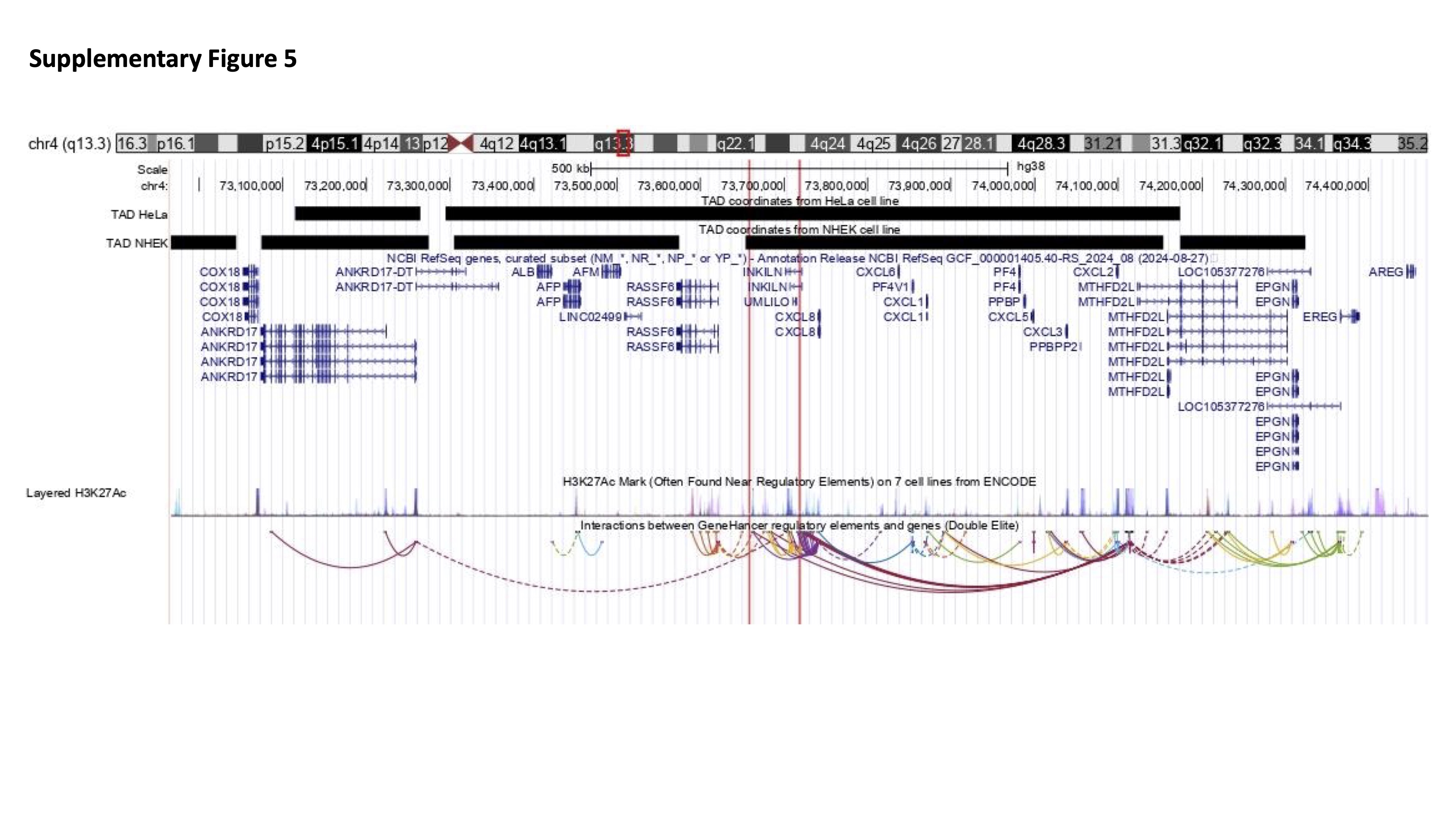

### Supplementary Figure 6

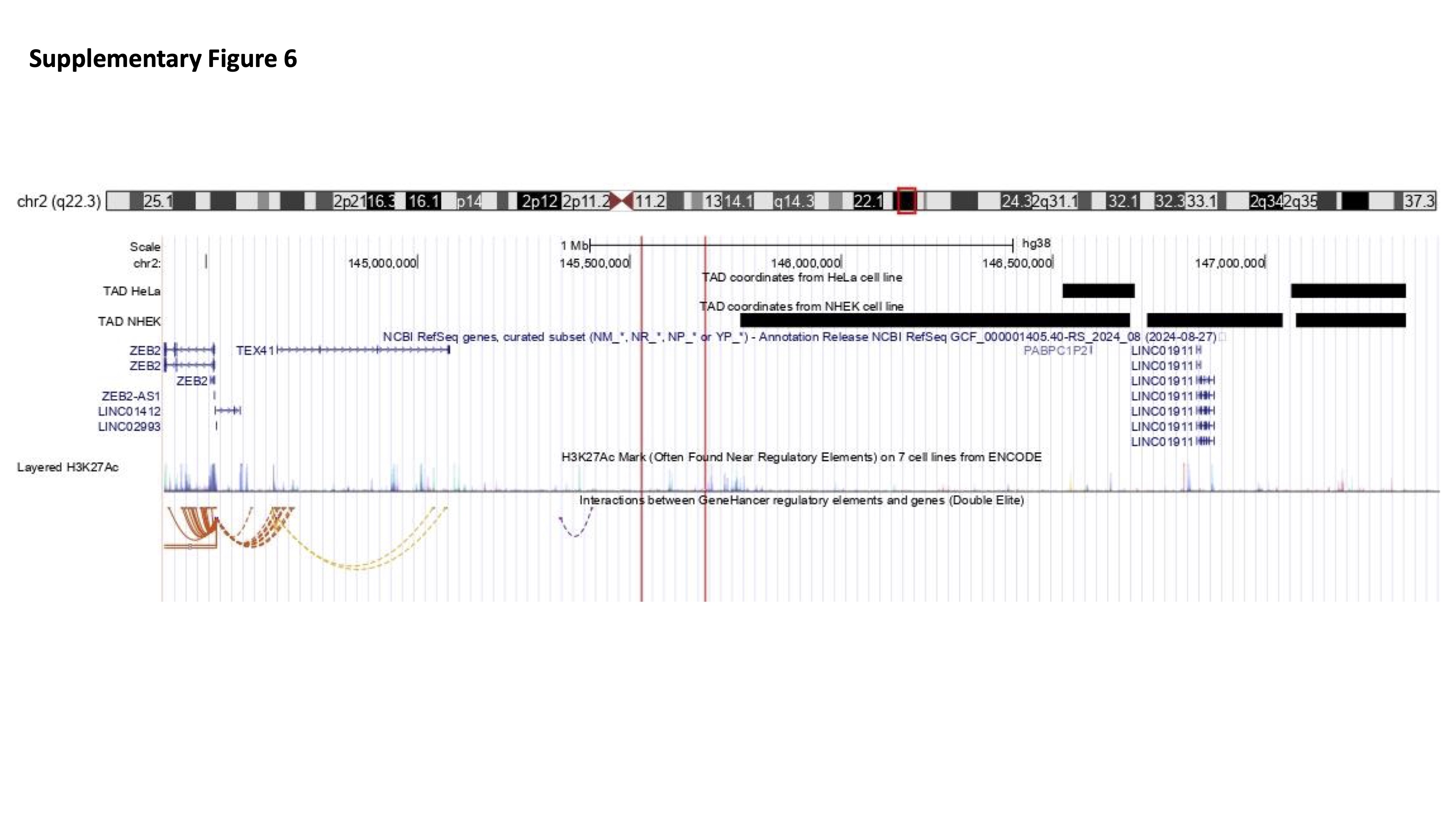
